# Characterizing Co-Circulating Respiratory Virus Genomic Diversity in Switzerland with Hybrid-Capture Sequencing and Phylogenetic Reconstructions: Insights into the 2023/24 Season

**DOI:** 10.1101/2025.07.21.664397

**Authors:** Charlyne Bürki, Matteo Carrara, Lea Fahrni, David Meyer, Anastasia Escher, Mara Neacsu, Selina Teufel, Christoph Noppen, Christiane Beckmann, Henriette Kurth, Franziska Singer, Louis du Plessis, Tanja Stadler

## Abstract

Respiratory viruses circulate yearly with strain-specific patterns. Although SARS-CoV-2 and Influenza A/B genomic surveillance is well-developed, most respiratory viruses are unevenly monitored, lacking geographical diversity to capture wider population dynamics. Consequently, insights into respiratory virus evolution are limited. We investigated the genetic diversity of these viruses within one country.

During the 2023/24 season, we conducted whole-genome sequencing of 1129 clinical samples using a hybrid-capture protocol. These samples were pre-tested by real-time PCR panels throughout Switzerland. Leveraging publicly-available full-length genomes, we constructed background datasets representative of geographical diversity and built phylogenies for all viruses with more than 40 new high-quality genomes from this study.

We detected 981 viruses and recovered 461 high-quality genomes, including 18 co-infections, from 437 PCR-positive and 6 PCR-negative samples. All viruses detected by PCR were also detected by sequencing in 56% of samples. The four most prevalent viruses were Influenza A/H1N1, SARS-CoV-2, RSV-A, and HPIV-3, and their seasonal spread was consistent with wastewater monitoring and influenza-like illness reports. Swiss viral genomes were representative of the global genomic diversity, with evidence for multiple introductions into Switzerland, and we identified putative Swiss clusters.

In this proof-of-concept study, we focus on 3 viruses (Influenza A/H1N1, RSV-A/B, and HPIV-3), and we demonstrate the streamlined implementation of a broad respiratory virus genomic surveillance workflow with an off-the-shelf protocol and publicly-available software. In addition, we highlight additional evolutionary insights that can only be derived from genomic surveillance. Going forward, this dataset will be a useful resource for future investigations into respiratory viral genomic diversity.

## Introduction

Accurately monitoring seasonal respiratory virus spread and evolution requires PCR testing, but genomic sequencing provides additional critical parameters relevant for public health authorities. For instance, sequencing allows for distinguishing local spread from imported cases [1,2], estimating epidemiological parameters during waves of co-circulating variants of a virus [3–6], tracking variants of concern [5,7,8], and monitoring genetic diversity associated with therapeutic escape mutations for a variety of infections [9,10]. These insights are particularly relevant when usual transmission patterns are disrupted, as observed during the unusual surge of respiratory syncytial virus A/B (RSV-A/B) cases in 2022 [11–13] and the “tripledemic” in 2023 [14,15].

A significant challenge in respiratory virus genomic surveillance is the limited geographical representation and the systematic under-representation of certain viruses in open-access databases. This disparity can lead to underestimating the true disease burden [16], particularly because limited localized studies often fail to capture the wider population dynamics of these rapidly evolving pathogens beyond the study population and period. As a consequence, there is limited knowledge about the evolution of most respiratory viruses, apart from SARS-CoV-2 and influenza viruses. Moreover, the interactions and competition between respiratory viruses remain to be clarified, particularly how the dynamics of one strain’s seasonal wave may impact that of another. Addressing these questions requires stable and broad longitudinal genomic surveillance programs, coordinated around the world.

To address the limited knowledge and evolutionary dynamics of additional respiratory viruses, surveillance efforts expanded beyond Influenza A/B and SARS-CoV-2, a development influenced by the disrupted circulation patterns observed during and after the COVID-19 pandemic [17–20]. Consequently, multiplex PCR panels, useful for simultaneously testing for several targets, are now often integrated in surveillance efforts, albeit with limitations in sensitivity. In parallel, respiratory virus *genomic* surveillance was improved through increased efforts in whole-genome sequencing. While more costly than amplicon-based methods, targeted enrichment by hybrid capture allows for efficient whole-genome sequencing of multiple pathogens in a single reaction with rapid turnaround times, yielding high-quality full genomes even in cases of co-infection [21–23]. In addition to these advances, the development of new bioinformatics tools and open-access databases has facilitated streamlined genomic analyses [24–26].

RSV-A/B genomic surveillance has benefited from this strengthened sequencing capacity. The virus is contracted by nearly all children in the first two years of life [27], and its subtypes A and B tend to exhibit a stable coexistence [28]. However, further studies are required, especially in light of the development of novel therapeutics, like the monoclonal antibody (mAB) Nirsevimab, targeting infants, and the RSV vaccine Arexvy, for adults [29,30]. Since July 2023, Nirsevimab was approved and used in the prevention of respiratory disease in infants less than 24 months of age in a few countries. First rolled out in Spain [31] and subsequently in the United States and Canada for the 2023/24 season [32], it has proven to be effective in reducing RSV-associated hospitalizations [29] by binding to the RSV fusion (F) protein, a protein key in viral penetration. In contrast, despite belonging to the same paramyxoviridae family as RSV-A/B and causing 13% of global childhood acute lower respiratory infections (ALRI) [33], Human Parainfluenza 3 (HPIV-3), also known as human respirovirus 3, has received considerably less global attention. It is reported as the third leading cause of pneumonia in young children [34], yet whole genome sequencing efforts remain sparse and geographically restricted [20,33].

Switzerland, with its three international airports and nearly 1.4 million (>15% of the population) daily train commuters [35] traveling on the densest rail network in the world [36], presents a highly mobile environment conducive to viral introductions. Aiming to characterize the co-circulating respiratory viruses in Switzerland, we collected and sequenced 1129 samples over the 2023/24 season, generating 461 high-quality genomes of 24 circulating respiratory viruses, and identifying 18 samples that were co-infected with multiple viruses. Incorporating publicly available full-length genomes representative of global diversity, we built phylogenies for all viruses with at least two high-quality genomes from this study, allowing us to identify epidemiological patterns of co-circulating viruses which exhibited localized, geographical clustering within Switzerland. We focus on the three most abundant viruses from this study (Influenza A/H1N1, RSV-A/B, HPIV-3), excluding SARS-CoV-2, and derive insights from full-length genome trees.

In our approach, we showcase the potential of characterizing the genomic diversity of co-circulating pathogens, anticipating that future studies will become more insightful as the amount of publicly available contextual full-length genomes increases. This study contributes majorly to increasing sequencing data in different virus families. Indeed, previous to this study, there was only one publicly available Swiss HPIV-3 full-length genome (Bioproject PRJNA73055) and one RSV-B genome dating from 2019 [37,38].

## Methods

### Sequencing

#### Collection, extraction, and sequencing

Over the 2023/2024 winter season, a total of 1129 archived nasopharyngeal swabs pre-tested by different real-time PCR panels (BIOFIRE® Respiratory 2.1 Panel, BioMerieux / Xpert® Xpress CoV2/Flu/RSV plus, Cepheid / Alinity m Resp-4-Plex assay, Abbott Molecular Diagnostics) for respiratory viruses of interest (RSV A/B, Influenza A (H1N1, H3N2) and B, SARS-CoV-2, Human parainfluenza 1-4, Seasonal coronaviruses (HKU1, NL63, OC43, 229E), adenoviruses, Rhino-/Entero-Virus, Metapneumovirus) were collected 5 to 14 days after testing. We first ran a pilot phase, collecting 94 samples from April 2023 to June 2023. Beginning in November 2023, we scaled up the collection, aiming to collect 94 samples every 2 weeks, randomly selecting 64 positive and 10 negative samples from the BIOFIRE® Respiratory 2.1 Panel, 10 positives from the Alinity m Resp-4-Plex assay, and 10 positives from the Xpert® Xpress CoV-2/Flu/RSV plus panel until mid-July 2024. The total bi-weekly number of samples collected was prone to variations depending on the availability of diagnostic samples. If not enough samples from a particular PCR panel were available, the remaining samples were collected from the other test panels from the same time period. A total of 1129 samples were sequenced, of which 128 tested negative by the associated PCR panel. Depending on the panel used, cycle threshold (CT) values were available.

Total nucleic acid was extracted from 400 µl of sample fluid using the Abbott mSample Preparation System DNA (Promega) on the Abbott m2000sp instrument. Library preparation was performed using the Illumina Respiratory Virus Enrichment Kit (https://emea.illumina.com/products/by-type/sequencing-kits/library-prep-kits/respiratory-virus-oligo-panel.html#tabs-9f8cdecf92-item-34e2792fef-overview) and sequencing was done on the NextSeq500 instrument. Additional metadata was collected including sampling location at the cantonal level, sampling date, PCR test result, and CT value, where available.

#### Bioinformatics,consensus genome assembly and analysis

Consensus genomes were generated using an in-house bioinformatics pipeline employing a two-step approach: (1) a metagenomic assessment of each sample to identify the appropriate reference sequences to map to and (2) read mapping and virus assignment based on the set of reference sequences identified in the first step. The rationale for the two-step approach is to complement the reference sequences used to construct the probes of the Illumina Respiratory Virus Enrichment Kit with references from untargeted viruses, since probes may also hybridise to related viruses not specifically targeted by the panel. The analysis pipeline was set up as a Snakemake (version 7.32.3) [39] workflow and is available as a Docker [40] image (https://hub.docker.com/r/ethnexus/revseq_bioinformatics_pipeline).

For the metagenomic assessment we used Kraken2 version 2.1.3 [41] with default settings and a custom database to obtain information on the metagenomic composition of each sample. For the custom database the organism definitions in the Kraken2 “Standard” database (https://genome-idx.s3.amazonaws.com/kraken/k2_standard_20240112.tar.gz, version January 2024) were adapted to reflect the overarching taxa diversity that are expected in the swab tests that were part of this study. Accordingly, we removed all archaea, plasmids, Univec_Core, bacteriophages, Poliovirus, as well as all plant viruses appearing in the database (https://www.dpvweb.net), all viruses responsible for Sexually Transmittable Infections appearing in the list at https://www.niaid.nih.gov/diseases-conditions/ sti-pathogens-and-syndromes, all Bovine Respiratory Syncytial viruses, and all Pandoraviruses. The hits detected in the metagenomic analysis were used to extend the Illumina reference panel, together with the human reference genome GRCh38 to allow the removal of host read contamination. Note that for Human Metapneumovirus (HMPV) we replaced the reference sequence from the Illumina reference panel with the sequence of Human Metapneumovirus isolate 00-1 (GenBank ID NC_039199.1), as the panel reference (GenBank ID NC_004148.2) was removed from the RefSeq database.

For the read mapping and subsequent virus assignment, for each sample raw sequencing reads were trimmed using Cutadapt version 4.9 [42], set to trim both primers and adapter sequences with options: *-b CTGTCTCTTATA -O 20 -m 20*. If adapter sequences were available in the metadata, we included them in the trimming by adding more *-b* options. Trimmed reads were aligned to the combined reference sequences using BWA mem version 0.7.17-r1188 [43] with default parameters.

In order to ensure that no host sequences remain for downstream processing, we performed a dehumanization step to remove all reads that mapped to the human reference. The remaining reads were further filtered using Samtools version 1.7 [44] to remove primary mappings and secondary mappings, as well as ambiguous mappings (i.e mappings that show an equally good alignment at multiple positions). The remaining virus-only, uniquely-mapped reads were further filtered using Picard [45] MarkDuplicates version 3.0.0 to remove optical duplicates.

Assignment of the infecting viruses present in a sample was performed in a two-step procedure using custom Python (version 3.11.9) scripts. In the first step, virus substrains were assigned as follows: for each reference virus substrain, we calculated the Reads per Kilobase per Million mapped reads (RPKM) to reflect how well it was mapped by the available reads. We computed the distribution of all RPKM values across the available references, and selected all substrains that show an RPKM value above the 95^th^ percentile as potentially infecting. This strategy can yield several substrain candidates per sample, e.g. in case of co-infections. In the second step, all reads from substrains part of the same strain are added together and the assignment procedure is repeated at strain level.

Consensus sequences were generated using BCFTools version 1.17 [44], including indel calling and normalization, and a threshold of 10 reads to mask low coverage positions. If more than one virus was identified as a potential infection, consensus genomes were assembled for all the likely identified viruses.

For downstream analysis, a virus detection was considered reliable if at least 10 uniquely mapped reads aligned to the reference strain genome. In other words, these are reads that did not map to any other reference genome. To overcome the variable quality yield of sequencing, we further selected sequences that had an unmasked consensus genome coverage of at least 20%, with a read depth of 10 (DP10 *>* 20%) and considered these as infections with high-quality genomes.

Each high-quality consensus sequence was assigned a random unique ID that was used to release the dehumanized raw data, the assembled consensus sequence, and associated metadata (sampling date, canton of sampling, PCR test result) to ENA under project PRJEB83635 (https://www.ebi.ac.uk/ena/browser/view/PRJEB83635?show=related-records). Additional resources were developed to facilitate data sharing, see Supplementary section “Project Database and UI”.

Subsequent analyses were performed in R, using packages dplyr [46], ggplot [47], and tidyr [48]. We retrieved GenBank files to annotate the genomes for the genome coverage plots. Weekly virus positivity rates were obtained from the publicly available national Swiss surveillance system (Sentinella) for the corresponding study dates (2023-03-15 to 2024-07-15) (https://www.idd.bag.admin.ch/topics/respiratory-pathogens/data). Wastewater 7-day median viral load estimates for Switzerland were obtained from the publicly available WISE dashboard (https://wise.ethz.ch/) for the same period.

The bioinformatics workflow is available on github at https://github.com/ETH-NEXUS/2023_cb_stadler_ReVSeq and as a docker image at https://hub.docker.com/r/ethnexus/revseq_bioinformatics_pipeline. The downstream analysis is available on github at https://github.com/charlynebuerki/swiss_co-circulating_viruses.

### Phylogenetic analysis

#### Context dataset construction

We assembled contextual datasets for all viruses with at least two high-quality genomes from this study and with at least 200 publicly available historical full-length genomes. Where enough sequences were available, we subsampled datasets to obtain virus-specific contextual datasets representative of temporal and spatial diversity. We briefly describe the context dataset construction for four viruses of interest below. The description for the other viruses can be found in the Supplement section “Other virus contextual dataset assembly”.

For both RSV-A and B, we adapted the RSV-A/B Nextstrain build (https://github.com/nextstrain/rsv) to query all publicly available consensus sequences, keeping only those with more than 10,000 non-N bases for the whole genome (equivalent to 65.8% or more of the genome coverage for RSV-A and 65.7% for RSV-B). Subsequently, we subsampled up to 300 genomes per country per year from 1925 until 2021, and up to 3000 genomes per country per year between January 2021 to August 2024, coinciding with the end of the sample collection period of this study.

For HPIV-3, contextual background sequences were obtained by querying consensus sequences under the associated taxon id (11216). The queried sequences were aligned and processed with Nextclade [25], where a coverage score was calculated for each consensus genome based on the same reference genome (accession reference: NC_001796) used to assemble the HPIV-3 Swiss samples of this study. We only kept sequences that were longer than 9,277 non-N bases (equivalent to 60% or more of genome coverage). We followed the same temporal and geographical subsampling as RSV-A and RSV-B.

For Influenza A/H1N1, we queried GenSpectrum (https://loculus.genspectrum.org/) to obtain all publicly available consensus sequences subtyped as both H1 and N1, subsequently randomly subsampling 30 sequences per year from Spring 2009 (2009-03-29), when the virus emerged, until 2022, and 300 sequences per country per year for the last 2 years. NA and HA segments were aligned, and sequences with segments containing more than 5% of unknown nucleotides were dropped.

The final numbers of genomes included in the datasets for each virus are shown in Table S1.

#### Phylogenetic reconstruction

For each virus, context genomes were combined with the core sequences of this project to build phylogenies representative of globally-circulating viruses. Maximum likelihood trees were reconstructed using IQ-TREE [49] using a general time reversible (GTR) substitution model for each virus. With TreeTime [50], tree branches were scaled to calendar time and ancestral states were inferred for each internal node following default parameters for Augur tree construction [24]. Amino acid substitutions were inferred and clades were assigned to nodes based on pre-existing clade annotations, if available, for the specific viruses. The HPIV-3 phylogeny was rooted to a sequence collected in 1957 in Washington, USA (NCBI accession: LC817393.1), given its high-quality, and an existing clade system presented by *Fahad N Almajhdi* [51] and refined by *Zhu et al.* [52] was incorporated. H1N1 HA and NA phylogenies were rooted to the pandemic strain (NCBI accession: CY121683.1) and the classic clade naming scheme available on Nextstrain (https://github.com/influenza-clade-nomenclature) was employed.

The phylogeny reconstructions are made in separate Nextstrain builds [24], using a reproducible snakemake workflow [39] available on the github repository of the downstream analysis: https://github.com/charlynebuerki/swiss_co-circulating_viruses/tree/main/Phylogenetic_analysis/Nextstrain_builds. In each build, the virus context dataset construction is included as part of the ingest module. All Nextstrain trees can be visualized with Auspice at https://nextstrain.org/community/cevo-public/ReVSeq-project.

We used ggtree [53] and custom scripts to adapt the Nextstrain trees to static images.

## Results

### Overview of circulating Swiss respiratory viruses

From the 1129 samples (N=1001 PCR-positive and N=128 PCR-negative) collected during the 2023/24 season from nasopharyngeal swabs pre-tested by PCR panels throughout Switzerland, we detected the presence of 981 viruses in 632 samples through sequencing with targeted enrichment by hybridization capture using the Illumina Respiratory Virus Enrichment Kit. Of those samples, 443 were of sufficiently high-quality to extract full-length genome sequences (Fig. 1 A). A comparison between PCR-positive sample detection and sequencing assignment at the strain level is detailed in Table S2.

**Figure 1:**
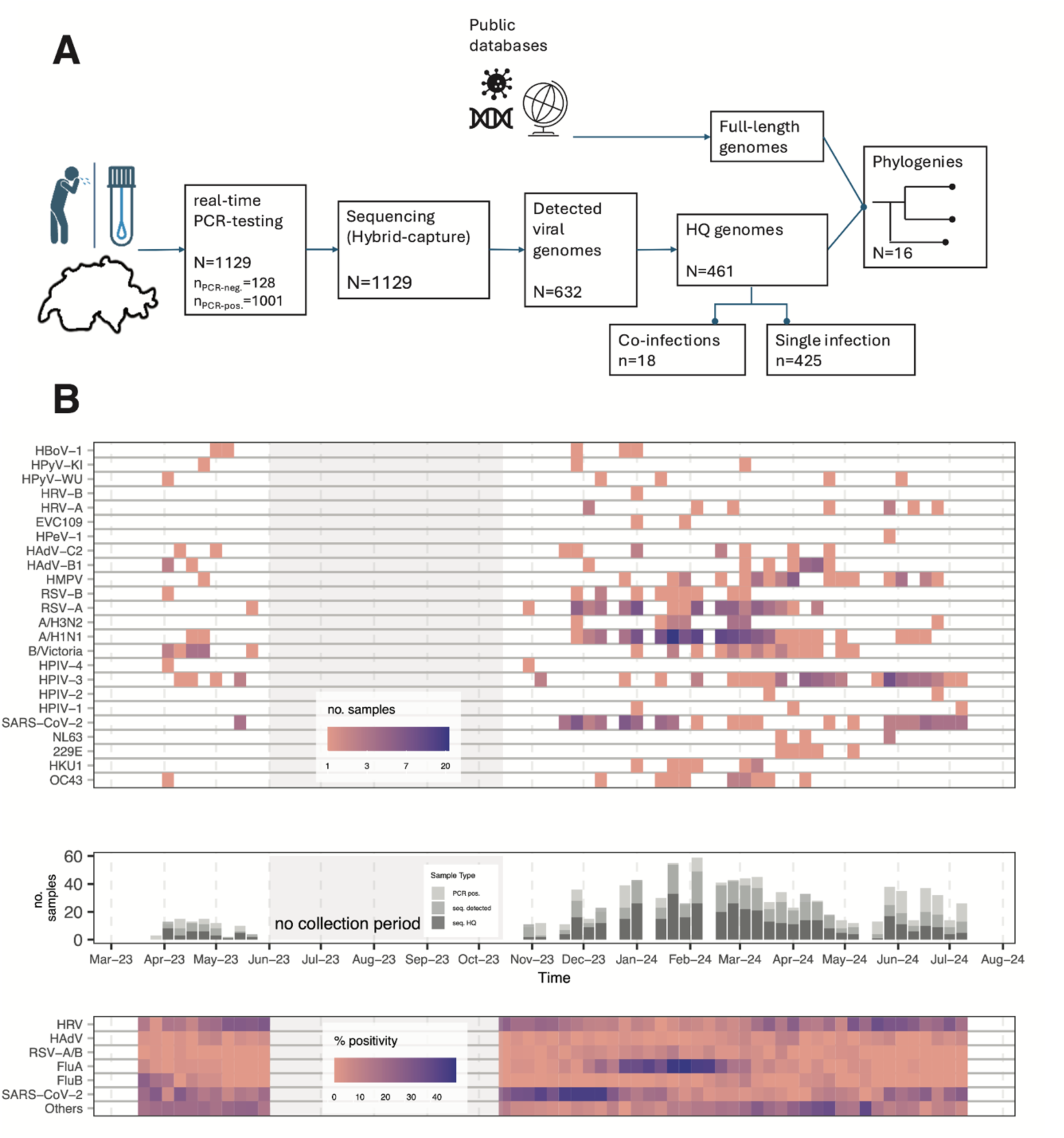
Study workflow and temporal overview of Swiss viruses circulating during the 2023/2024 season: **A.** The study workflow with intermediate steps and number of samples at each step of the study. The workflow spans from the collection of samples during the study period to the reconstruction of phylogenetic trees made from this study’s samples incorporated with contextual datasets for each virus with at least 2 high-quality sequences from this study (N=16). **B.** (top) The temporal overview of all high-quality genomes (N=461, DP10>0.2) obtained in this study, classified by virus, aggregated at a weekly resolution, where darker colors indicate greater numbers of high-quality genomes obtained during a week. (middle) The relative positivity rate for high-quality samples at a weekly frequency (dark grey= number of high-quality samples, grey= number of detected samples by sequencing, light grey= number of PCR-positive samples). (bottom) The reported, publicly available national surveillance system (Sentinella) weekly positivity rate for 7 categories of pathogen from PCR-positive samples sampled in Switzerland, where a darker color indicates a greater positivity rate (for a more detailed comparison with samples from this study, see Fig. S1 and Table S3).

High-quality genomes consisted primarily of five viruses (320 of 461): Influenza A/H1N1 (N=118), SARS-CoV-2 (N=61), RSV-A (N=56), HPIV-3 (N=56), and HMPV (N=29). The pilot phase ran from April 2023 to June 2023, and the main collection period, where we collected 94 samples every two weeks, ran from November 2023 to mid-July 2024, with slight fluctuations depending on the availability of diagnostic samples. Positive PCR tests, virus detections, and high-quality genomes peaked between January and March 2024. The bulk of infections detected from November 2023 to mid-January 2024 consisted mainly of SARS-CoV-2 infections. From mid-December 2023 to the beginning of April 2024, we observed a high level of Influenza A/H1N1 and RSV-A infections, both detected by PCR positivity tests and through sequencing, with the Influenza A/H1N1 infections reaching a peak in mid-January 2024, slightly earlier than the RSV-A peak (see Fig. 1 B). RSV-A samples represented 83.6 % (56/67) of high-quality RSV-A/B genomes. Similarly, the majority of Influenza virus detections were of Influenza A/H1N1, with only a low number of Influenza A/H3N2 (between December and July 2024) and Influenza B (in April 2023 and again in March and April 2024) detections.

High-quality HMPV genomes were recovered from samples between March and April 2024 and the bulk of HPIV-3 high-quality genomes occurred in June 2024. Other types of high-quality parainfluenza virus genomes were assembled from samples distributed sporadically throughout the study period. High-quality seasonal coronavirus genomes were recovered from samples between December 2023 and June 2024, with earlier occurrences of coronavirus OC43 and HKU1 (mid-December to mid-March). High-quality Adenovirus genomes (C2 and B1) were assembled throughout the winter season, and other viruses such as rhinovirus A, enterovirus C, and polyomavirus were found sporadically during the study period.

While our sampling was independent of the incidence of any particular pathogen (but instead we aimed to collect a fixed number of samples every two weeks), virus collections were nevertheless representative of the Swiss situation during the 2023/24 season, as evidenced by reasonably high positive correlations between the detected strains and the two national surveillance systems (Sentinella and wastewater monitoring), for available viruses (Table S3). For instance, the *R*^2^ between Influenza A detections from sequencing and wastewater monitoring was 0.71 and 0.60 for Sentinella surveillance. Furthermore, the peaks of influenza A and SARS-CoV-2 from the detected viruses identified by sequencing coincided with the peaks detected in the Swiss national Sentinella and wastewater surveillance systems (see Fig. S1 and Fig. S2). While the Sentinella system did not detect a well-pronounced wave of RSV-A/B, the study samples and wastewater viral load estimates are correlated and reveal a wave characterized by a larger spread over the winter than the influenza A/H1N1 peak (see Table S3 for all correlation values).

### Sequencing results

#### Concordance of PCR tests and sequencing results

When treating PCR-testing as the ground-truth, concordance between the strains identified by PCR testing and those detected by sequencing was consistent (every strain detected by PCR was also detected by sequencing, and additional viruses could also be detected by sequencing) or partially consistent (there is at least one match between PCR and sequencing detections, but additional viruses are detected by PCR) for 56.2% (634/1129) of samples (see Fig. 2 A). However, among the 1001 samples that tested PCR-positive for one or more viruses, no virus infections could be confirmed by sequencing in a third of these (n=383). We examine the possible causes for this lack of sensitivity in the discussion.

**Figure 2:**
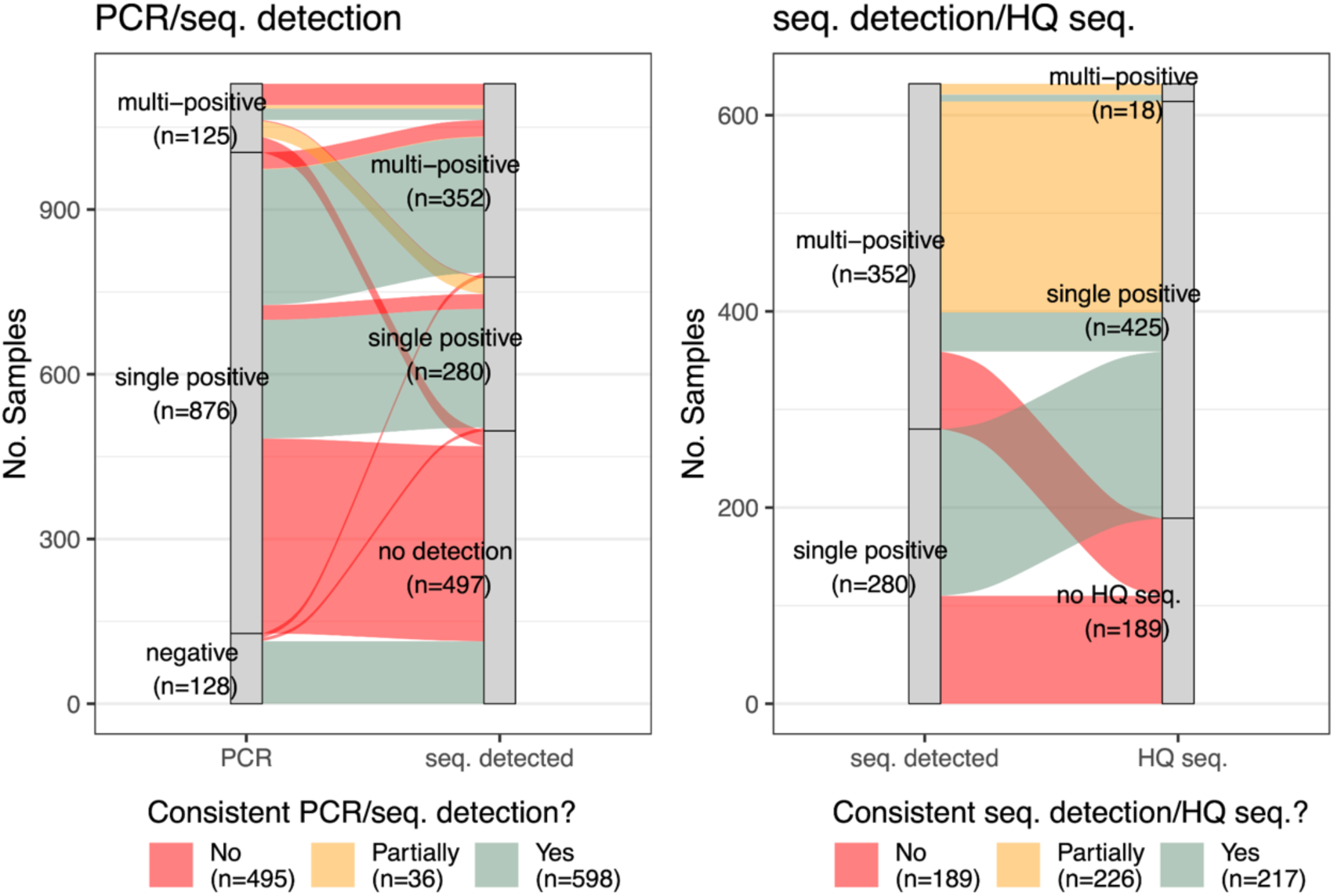
Consistency of strains identified by testing method on all samples between. (left) PCR test and detected strains by sequencing and (right) strains detected by sequencing and high-quality genomes. A consistent link (green) implies that the same strain is present in both testing methods, and in the case of a single to multi-positive match, it includes additional strains not identified in the single infection. A partially-consistent match (orange) implies that only a subset of the identified strains are identified or recovered. An inconsistent result (red) implies a total mismatch between the two.

We found that among samples where sequencing identified a unique positive infection, sequencing results perfectly matched the PCR result 77% of the time (216/280). Partially consistent matches represented 10.4% (29/280) of single-positive sequencing detections, while the remainder (n=35) were mismatches. In 18 out of the 29 partially-consistent matches, samples tested PCR-positive for rhino/enterovirus, but these viruses were not detected by sequencing. Similarly, 14 out of the 35 mismatches were identified as PCR-positive for rhino-/enterovirus infection, however we detected an infection by a virus from a different family.

Among samples where sequencing detected a co-infection with more than one virus (multi-positives hereafter), sequencing results were consistent with the corresponding PCR test result in 76% of samples (268/352), and in 40% (106/268) of these, we detected additional substrain(s) not identified by PCR. The remainder of multi-positives detected consisted of 7 samples partially matching to their PCR test and 77 with entirely inconsistent matches.

Overall, we detected a virus not included in any of the PCR panels in 145 samples. Among the 495 inconsistent matches, 251 (50.7%) either tested PCR-positive for only rhino-/enterovirus (n=210) or that virus constituted at least one of the multi-PCR-positive strains (n=41). Interestingly, we detected viruses in 14 PCR-negative samples, subsequently leading to 7 high-quality genomes, consisting of 4 RSV-A infections, including a co-infection with an enterovirus, 1 RSV-B, and 1 Influenza A/H1N1 infection. Since RSV and Influenza targets are included in all PCR panels used (Table S2), this illustrates the ability of hybrid capture sequencing to not only detect, but also recover high-quality genomes from samples that tested PCR-negative.

From the 632 samples with virus detections by sequencing, resulting in 981 detected viruses, we generated 461 high-quality genomes, including 18 co-infections. In the 49.4% (210/425) of cases where only a single, high-quality genome was recovered, it was consistent with the detected strain (see Fig. 2 B). Of these, 40 samples were detected as a multi-positive infection with substrains of the same virus (e.g. RSV-A and RSV-B), but a high-quality genome could be assembled for only one of the substrains. The remainder of the samples (n=215) from which only a single high-quality genome could be generated were from samples detected as multi-positive infections, with viruses from different families, where target-enrichment resulted in enough reads to allow assembling a high-quality consensus genome for only one of the viruses. In the case of the 18 high-quality co-infecting genomes, we found that seven were in exact concordance with the multi-positive detection. We were unable to assemble a high-quality genome for 30% (189/632) of the infections detected by sequencing, and adenovirus and entero/rhinovirus infections accounted for 44% of these (84/189).

#### Identification of co-infections

We generated high-quality full-length genomes from 18 (4.1% of high-quality genomes) co-infections (see Fig. 3), less than previous clinical respiratory co-infection estimates, ranging from 10 to 20% [54], with more frequent prevalence rates reported in pediatric cases [55,56]. RSV-A was the most common (n=7), representing 12.5% (7/56) of all high-quality RSV-A sequences in our dataset. We also generated 4 Influenza A/H1N1 genomes from co-infections (with RSV-A, HMPV, polyomavirus and SARS-CoV-2), representing 3.4% of all Influenza A/H1N1 high-quality genomes (4/118 samples). The Influenza A/H1N1 and SARS-CoV-2 co-infection was the only SARS-CoV-2 co-infection in our dataset for which we could generate a high-quality genome. Polyomaviruses (KI Polyoma and WU Polyoma viruses), viruses not covered by the PCR panels, were found with a broad range of co-infecting viruses, accounting for 22% (4/18) of all co-infections. This virus was also detected in four samples without any other co-infecting viruses.

**Figure 3:**
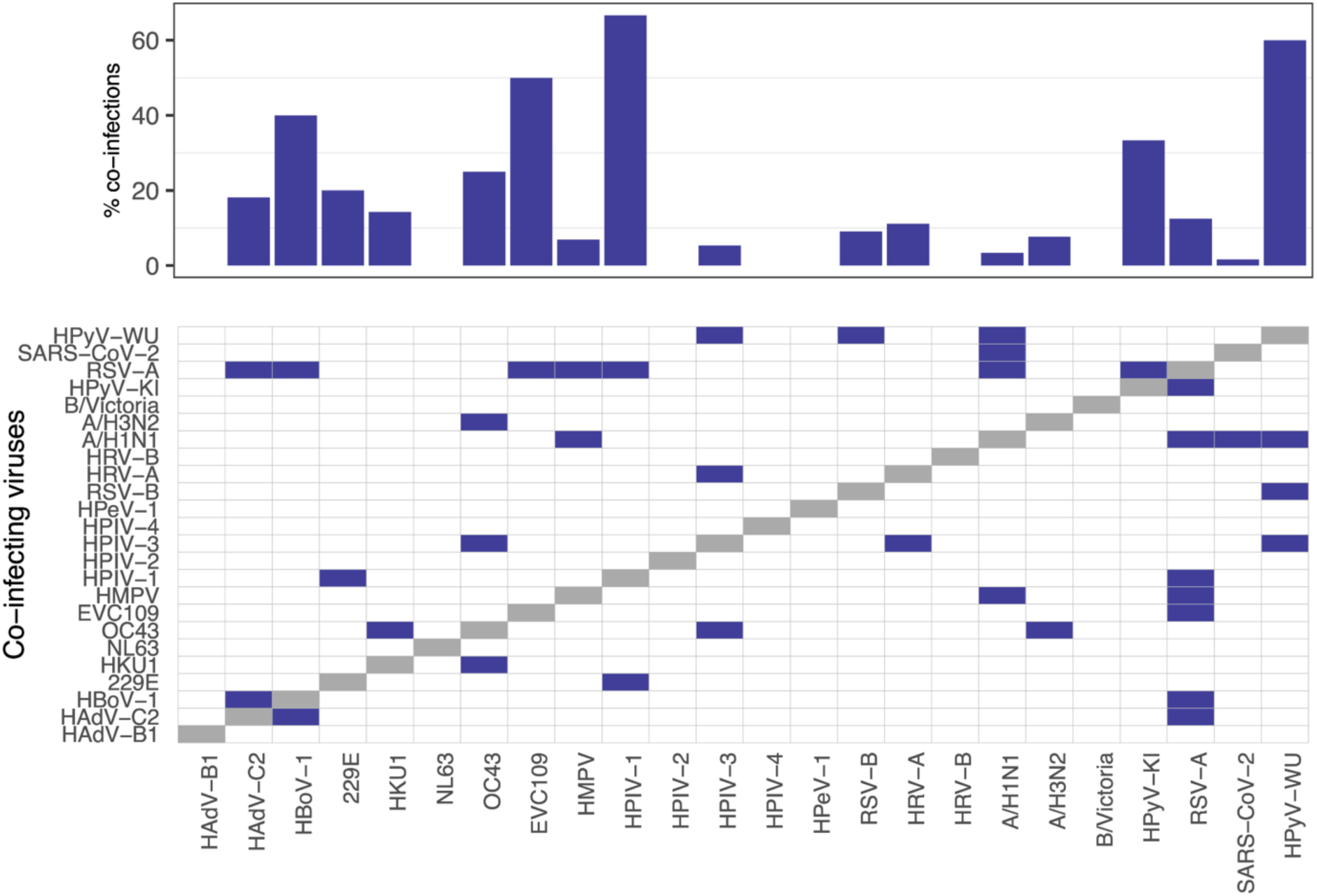
Co-infections detected in high-quality genomes: (top) per-virus strain co-infection rate detected in all high-quality genomes (bottom) the co-infection matrix: an entry indicates a co-infection between a virus in a row/column combination.

#### Genome coverage

In the infections where we could extract a high-quality full-length genome (hereafter high-quality genome), the depth coverage varied between viruses and samples; however, the intra-sample depth coverage variability was low. We highlight examples of negative single-stranded RNA viruses (ssRNA (-)).

HPIV-3 (n=56 genomes) had a median coverage across the whole CDS above 100X, including the F gene, although the non-coding region located between the M and F genes had consistent drop-out in all genomes. All HMPV samples collected between February 29^th^ to May 21^st^ were characterized by across-the-genome, above 100X coverage with a notable drop-out on the G gene; however, samples collected outside this period were characterized by notable drop-outs at the beginning of the P and F genes and throughout the S and G genes. On the other hand, RSV-A and B genomes yielded consistent >100X coverage in a majority of samples, without any systematic dropouts (See Fig. 4 for RSV-A and HPIV-3 coverage and Fig. S3 A and B for RSV-B and HMPV coverage). Other parainfluenza viruses (1 and 2) collected in this study had consistent high coverage but our dataset contained only a small number of these viruses.

**Figure 4:**
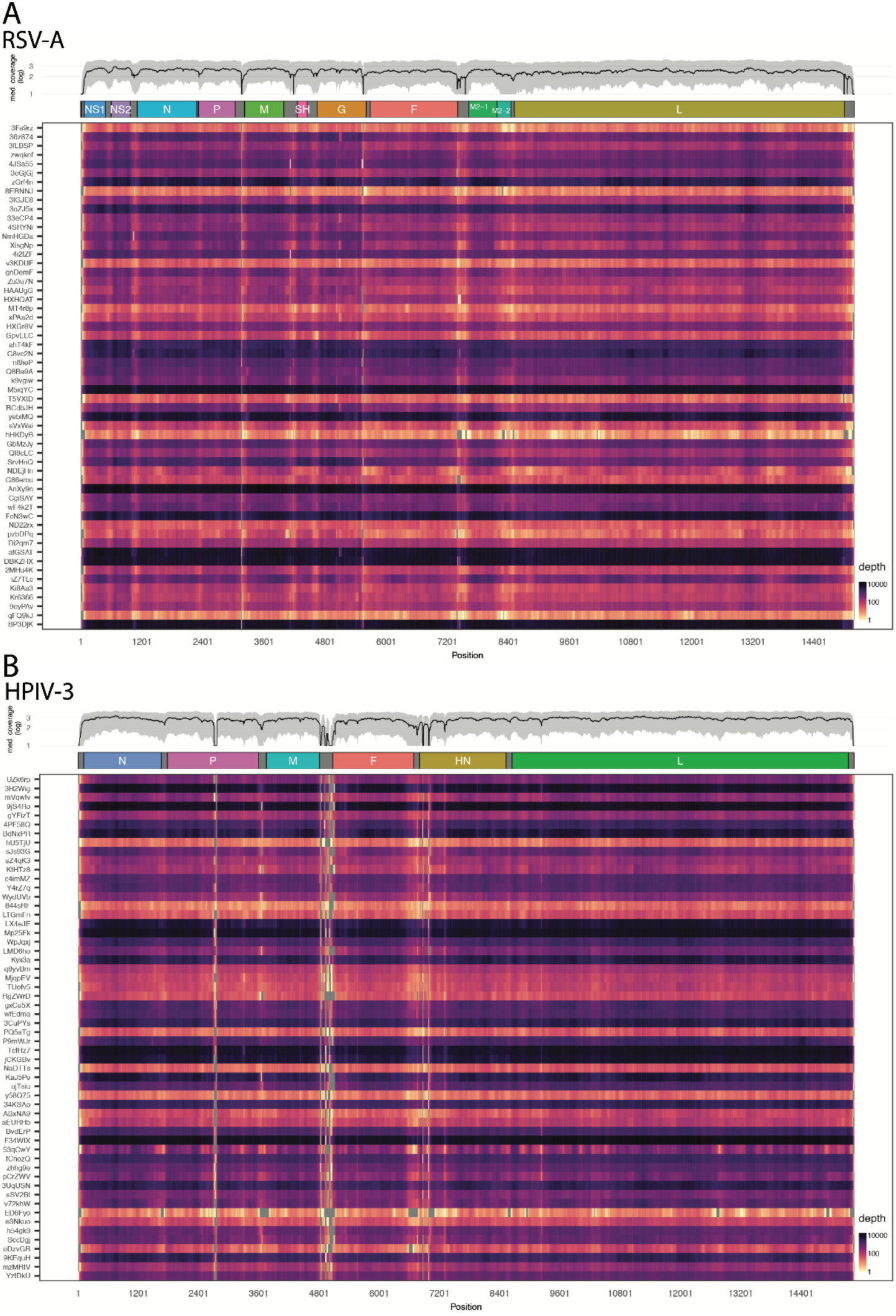
Whole-genome coverage of RSV-A and HPIV-3: The 56 RSV-A in **A.** and 56 HPIV-3 in **B.** annotated, whole-genome coverage of high-quality genomes obtained in the study. The median coverage in black and the 10*^th^* to 90*^th^* quartile in shaded grey calculated at each basepair is plotted in the top panel, and the individual genomes’s depths are plotted as a heatmap below, where a darker color indicates greater depth. Grey indicates a depth coverage of 0. Samples are ordered in increasing collection date order.

In the Orthomyxoviridae family, the 118 Influenza A/H1N1 samples had a median coverage at each base pair well over 100X, on each of the 8 segments. However, there were some samples with lower coverage throughout the season, with no discernible pattern to explain this drop in coverage (see Fig. S3 C for the HA and NA segments). Influenza A/H3N2 and Influenza B/Victoria followed the same pattern as Influenza A/H1N1, but our dataset does not contain enough samples to draw any conclusions.

The 5 types of viruses included in the coronaviridae family – SARS-CoV-2, seasonal coronavirus HKU1, NL63, OC43, and 229E – were all well-covered, especially on the spike protein, although we found a drop out in the HE and Spike protein in 3 out of 6 high-quality HKU1 genomes. All coverage plots can be found at https://github.com/charlynebuerki/swiss_co-circulating_viruses/blob/main/coverage_plots_all.md.

### Phylogenetic analysis

We assembled contextual datasets for all viruses with at least 200 publicly available historical full-length genomes and with at least 2 high-quality genomes from this study to construct ML phylogenetic trees. These viruses are RSV-A and RSV-B, Influenza (A/H1N1, A/H3N2, B/Victoria), HPIV (1,2,3), HMPV, SARS-CoV-2, seasonal coronaviruses (OC43, HKU1, NL63 229E), and adenoviruses (B1, C2). The corresponding trees can be explored in Nextstrain: https://nextstrain.org/community/cevo-public/ReVSeq-project. Below, we only discuss phylogenetic trees with more than 200 publicly available full-length genomes containing at least 40 high-quality genomes from this study. Although we obtained 61 SARS-CoV-2 samples from this study, we do not discuss the phylogeny and refer the reader to the available Nextstrain tree, as SARS-CoV-2 is actively monitored by the Federal Office of Public Health (FOPH) of Switzerland. All other viruses from this study did not meet the above criteria.

#### RSV-A/B

The RSV-A phylogeny consisted of 3’074 tips: 1’633 from North America, 1’032 from Europe (including 56 Swiss sequences), 147 from Asia, 141 from Africa, 41 from South America, and 79 from Oceania. All 56 Swiss sequences in the tree were generated in this study. The sequences spread between 8 different clades (A.D.1, A.D.1.5, A.D.1.6, A.D.2.1, A.D.3, A.D.3.1, A.D.5.1, and A.D.5.2), with 18 samples adhering within the A.D.5.1 and A.D.5.2 clades (top inset of Fig. 5 A.), 12 to A.D.3 and A.D.3.1 clades (middle inset of Fig. 5 A), and 25 samples clustering between the A.D.1, A.D.1.5 and A.D.1.6 clades (bottom inset of Fig. 5 A). A.D.1, A.D.5.1 and A.D.3 made up the majority of the clades, representing, respectively, 44.64%, 32%, and 21.4% of Swiss samples. Although A.D.1 was made up of geographically diverse cantonal samples (Northwestern Switzerland, Vaud/Geneva, and Eastern Switzerland), the A.D.3 and A.D.5 clusters contained more Northwestern Swiss samples.

**Figure 5:**
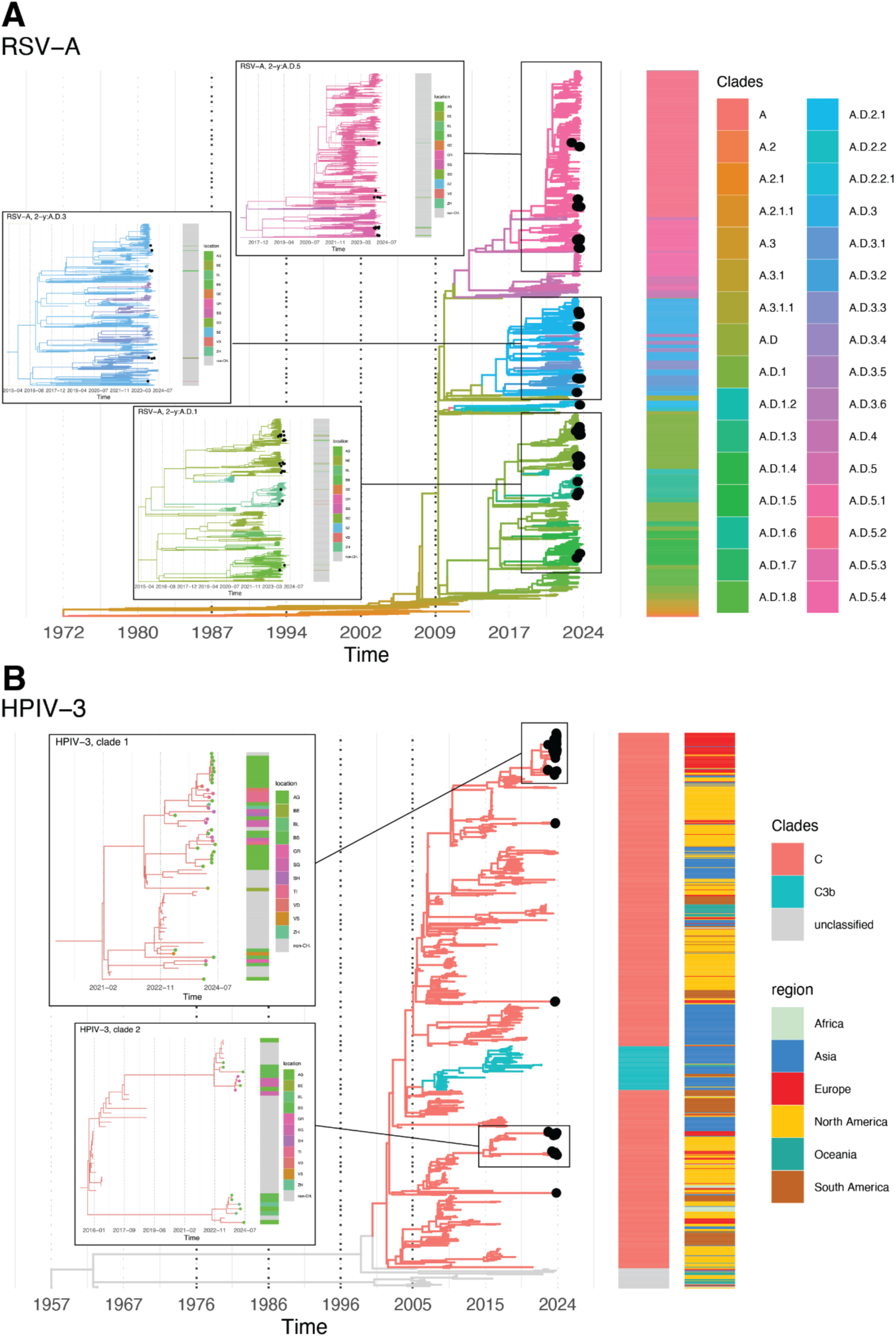
Whole-genome phylogenies of RSV-A and HPIV-3. The maximum likelihood phylogenetic time trees generated with the Nextstrain builds color-coded by clade and labeled with black dots for this study’s samples. in **A.**, the RSV-A tree with 3 insets into clade A.D.5 (top; pink), A.D.3 (middle; blue) and A.D.1 (bottom; green), with the cantonal location of this study’s samples to the right of the insets. The clade labeling is from Nextstrain. In **B.**, the HPIV-3 phylogenetic tree with additional labeling by geographic region; the two insets are clade 1 (top), with a majority of this study’s samples originating from Northwestern Switzerland and clade 2 (bottom), where there is more diversity in the samples’ location. The clade labeling is taken from [52].

While the RSV-A Swiss samples spanned the diversity of circulating clades, RSV-B Swiss samples of this study spread across one large clade: B.D.E.1 and its subclades B.D.E.1.2 and B.D.E.1.4 (see Fig. S4) while the unique ancestral Swiss sequence, dating from January 2019, was part of the B.D.4.1.1 clade. The large B.D.E.1 clade also contained the majority of tips included in the phylogeny (1’895 out of 2’643 tips). The Swiss RSV-B samples from this study were defined by previously reported homoplasic mutations on the G gene [19]: I252T, I268T, S275P, with the additional new Y285H mutation. An F gene mutation, S389P, was also observed, which has been reported in multiple countries.

We also screened the F genomes for escape mutations of both strains predicted to be in the antigen binding site of Nirsevimab [9,19,57] but found no predicted mutations in Swiss samples.

Finally, we calculated the patristic distance of Swiss tips from the mutation trees of RSV-A and RSV-B and found that the median (IQR) patristic distance of Swiss RSV-A tips (0.0191 (0.01040-0.0205)) was higher than the median (IQR) patristic distance of Swiss RSV-B tips (0.0066 (0.00515-0.0072)) and their distributions differed significantly (Kolmogorov-Smirnov Test D=0.884; p-value < 0.001).

#### HPIV-3

In this study, we generated 56 high-quality HPIV-3 genomes, plus 7 other parainfluenza virus genomes (3 HPIV-1 and 2 HPIV-2, 2 HPIV-4). In addition to the HPIV-3 Swiss dataset created in this study, another full-length Swiss genome of HPIV-3 dating from May 2011 was publicly available and included in the phylogeny.

The HPIV-3 phylogeny was made up of 823 tips in total: 352 from North America, 210 from Asia, 112 from Europe (including the 56 Swiss samples of this study and the one historical Swiss sample), 98 from South America, 33 from Oceania, and 18 from Africa. Samples from this study mainly aggregated in two large clades in the tree: clade 1 (top inset of Fig. 5 B), and clade 2 (bottom inset of Fig. 5B). Clade 1 was principally made up of Swiss samples, with samples originating from across Switzerland, primarily Southern (Ticino, Valais, and Vaud) and Northwestern Switzerland (Aargau, Basel-Land, Basel-Stadt). Non-Swiss samples in the clade were nested within the genetic diversity of Swiss samples, with the majority originating from France. Most of the samples from the canton of Aargau clustered in two closely-related subclades of 4 and 9 samples each and were collected between May 21^st^ and June 17^th^ 2024, with the 9-sample subclade exhibiting a ladder-like phylogeny. Within clade 2, Swiss samples clustered in two genetically distinct subclades, each attached by a long branch to a background of older sequences from the USA. Swiss samples originated from across Northern Switzerland (Cantons Basel-Stadt, Basel-Land, Zürich, St-Gallen, and Aargau) and one of the Swiss subclades contained 5 closely-related Japanese sequences. Interestingly, 2 of the 3 samples from the canton of Bern were identical and attached to a long branch stemming from a clade of Peruvian samples dating from 2012. The Swiss sequences falling into clusters does not give hints to local transmission chains though, as there are 56 Swiss sequences and only 31 contemporaneous contextual sequences. Thus, Swiss sequences cluster due to their outsized representation in the tree and more global sequences are required to assess whether the Swiss sequences cluster and represent putative Swiss transmission chains.

#### Influenza A/H1N1

The Influenza A/H1N1 HA phylogeny consisted of 1607 tips: 1’164 from North America, 10 from Oceania, 100 from Asia, 11 from South America, 8 from Africa, and 314 from Europe (including 153 Swiss samples: 110 from this study and 43 from publicly-available genomes). The 110 A/H1N1 HA genomes spread between two clades: 5a.2a.1 and 5a.2a (see Fig. S5). Within clade 5a.2a.1, the Swiss samples from this study were characterized by short branches, each attached to a background of USA sequences from the previous fall, with nearly identical HA genomes. These samples spanned the diversity of the whole clade. Half of Clade 5a.2a was made up of Swiss samples from this study and other surveillance sources. Samples from this study covered all regions of Switzerland. Non-Swiss sequences were mainly from the USA, with other European sequences nested within the genetic diversity of Swiss samples. In two instances, we observed a cluster of identical genomes during the Influenza A/H1N1 wave (mid-January to mid-March 2024): in one case, 20 identical Swiss sequences from Northwestern Switzerland (Cantons Basel-Land, Basel-Stadt, Schaffhausen, and Aargau) (see inset of Fig. S5), and 13 identical genomes from Switzerland, the UK and the USA. The remainder of the genomes attached with short branches to a clade consisting of Swiss samples, mixed in with geographically-diverse sequences.

## Discussion

We set out to characterize the genetic diversity of respiratory viruses circulating during the 2023/24 season in Switzerland using an off-the-shelf hybrid capture protocol. We extracted high-quality genomes of 461 viruses, including 18 co-infections. The viruses characterized were representative of the Swiss situation with respect to Influenza-like illnesses, as evidenced by high correlations with two surveillance systems (Sentinella and wastewater monitoring). Focusing on respiratory viruses with at least 40 high-quality genomes (RSV-A, HPIV-3, Influenza A/H1N1) and a large background dataset (N>200 full genome sequences), excluding SARS-CoV-2, we investigated the transmission patterns based on phylogenies on a local scale, serving as valuable public resources for future respiratory pathogen genomic studies. With the exceptions of SARS-CoV-2 and influenza viruses (A/H1N1, A/H3N2, B/Victoria), prior to this study, public repositories contained at most one, or zero, Swiss full-length genomes for studied respiratory viruses. For HPIV-3 alone, despite not specifically targeting sample selection for this virus, we generated 56 high-quality genomes from Switzerland, while only one sequence was previously available. These genomes contribute to an increase in over 6% of all publicly available HPIV-3 genomes, underscoring the underrepresentation of this virus in genomic databases and demonstrating the ease with which we can rapidly augment our knowledge using hybrid capture panels (refer to Table S4 for a list of publicly available full genomes of the different respiratory viruses discussed).

Employing a hybrid-capture method for respiratory virus surveillance instead of PCR testing could prove useful to obtain more precise information on the genomic diversity of circulating viruses, namely, for subtyping strains (differentiating between RSV A and B, or Influenza A/H1N1 and A/H3N2), for covering a broader range of pathogens than those included in commonly used PCR panels, such as polyomaviruses, and for identifying mutations of interest with a short turnaround time. At scale, the per-sample reagent costs of hybrid-capture sequencing are comparable to those used in multiplex PCR testing. However, in case timely results are required, PCR-testing remains a necessary component of genomic surveillance. In our study, sequencing in general exhibited lower sensitivity than PCR-testing (see Limitations and future directions), however in some cases allowed detecting infections and even recovering high-quality genomes from samples that tested PCR-negative for exactly those viruses. Omitting samples testing PCR-positive for rhino/enteroviruses, we consistently detected or partially detected by sequencing the same PCR-positive virus 66% (449/679) of the time, in addition to detecting strains in 14 of the 128 PCR-negative samples. Furthermore, we obtain high-quality full genomes for 65.2% (443/679) of the non-rhino/enterovirus PCR-positive samples. Going forward, with an envisioned increase in sensitivity of hybrid capture methods, sequencing could yield results comparable to PCR-testing, with the added benefit of producing high-quality genomes, allowing for a finer-grained overview of the respiratory virus landscape and facilitating downstream phylogenetic and phylodynamic analyses.

In what follows, we discuss the observations derived from the phylogenies highlighted in this study.

### RSV-A/B co-circulated with higher prevalence of RSV-A, with no notable fusion gene mutations reported

RSV A and B appeared to be mixing fast globally, resulting in Swiss samples representing global RSV-A/B genomic diversity in the surveilled season. RSV-A has increased in its global circulation since 2022 [58–60] and this was reflected in our Swiss dataset (83.6 % (56/67) of RSV samples were RSV-A), also resulting in some Swiss RSV-A clusters which might indicate local transmission clusters. The statistically significant higher patristic distance of Swiss RSV-A tips relative to RSV-B, indicates a higher genetic diversity, and suggests that the 2023/2024 RSV-A wave was characterized by long-term cocirculation of multiple genetically distant clades that originated more than 10 years ago.

For RSV-A in particular, we reported samples spanning 9 different clades, with samples aggregating mainly within 3 clades: A.D.1 (44.64%), A.D.5.1 (32%), and A.D.3 (21.4%), as other recent studies have reported in the USA and Canada [59,61]. Further, the phylogenetic analysis of RSV-A showed evidence of localized expansion. From the 12 Swiss samples clustering to A.D.3 and its subclade A.D.3.1, 8 samples formed 2 monophyletic clades consisting solely of Swiss sequences: 4 samples from Northwestern Switzerland expanded from a Swiss TMRCA inferred to October 2021, characterized by 2 homoplasic G gene mutation: L71P and T200A, and the 3 others were from two cantons: Basel-Stadt and St-Gallen. We also found evidence for local expansion of a single Swiss variant in the A.D.5 and A.D.1 clades. For example, in A.D.1, 5 samples collected from February 17^th^ to March 30^th^, formed a monophyletic Swiss clade, and were uniquely characterized by 5 homoplasic G gene mutations (E271X, E295X, Y297N, H304Y, *322X). While the high genetic similarity, close sampling times, and geographical origins are consistent with localized transmission, the clustering of Swiss samples cannot be directly interpreted as Swiss transmission chains given the low number of Swiss samples in the tree (at the time of writing these were the only publicly available Swiss RSV-A genomes).

In comparison, all Swiss RSV-B samples aggregated within a monophyletic clade, B.D.E.1, like studies focused on other geographic regions spanning the 2022/23 and 2023/24 winter seasons reported [57–60], despite a background dataset as large and geographically diverse as RSV-A. However, the Swiss samples are spread throughout the clade, suggesting continual re-introduction into Switzerland. Although some samples within the B.D.E.1 clade formed distinct clusters in its subclades B.D.E.1.2 and B.D.E.1.4, we found almost no clusters originating from a single Swiss sample.

We did not observe any Nirsevimab escape mutations in our RSV A or B high-quality genomes. In line with other studies reporting no widespread increases of escape mutations over the last two years globally, imports into Switzerland should be infrequent. As for the situation within Switzerland, evolution and spread is not yet likely as Nirsevimab was only largely deployed to a specific target group in Switzerland during Fall 2024, after this study’s sampling period.

The collected Swiss RSV dataset is crucial to have as a background to investigate future evolution and spread of escape mutations as the mAB is deployed more widely. Moreover, the dataset allows to monitor the impact on RSV evolution of the planned deployment of the new adult vaccine (Arexvy) in Switzerland in the coming seasons.

### HPIV-3 circulation was characterized by potential localized, within-country spread

Human Parainfluenza infections reported in this study were primarily from HPIV-3, with sampling occurring predominantly in late Spring. The HPIV-3 phylogeny was made up of four currently circulating clades (more recent than January 2022) and a few distinct long branches. Swiss samples alone spanned two of the four clades and additionally formed three long branches in the tree. Notably, phylogenetic analysis revealed that 68% (38/56) of the HPIV-3 high-quality genomes from this study clustered into a distinct group primarily composed of Swiss sequences (38/63), although it also included representatives from France, the USA, Japan, and Thailand. That cluster was characterized by a few mutations, notably two mutations in the HN and F genes: *573L in the HN gene and I521T in the F gene. Within the Swiss samples of this cluster, we observed potential evidence of localized spread. For example, nine genomes originating from a single canton, Aargau, formed a sub-cluster characterized by an additional F gene mutation (I499V), suggesting expansion from a single local variant. This regional pattern extended to other cantons represented in the phylogeny, whether from the Bern samples uniquely clustering together on a long branch or the close proximity of sequences from the neighboring cantons of Basel-Stadt and Basel-Land. The only other publicly available full-length Swiss HPIV-3 genome, which dates back to 2011, was found to be more ancestral to clade 2 identified in our study.

We emphasize that these clustering patterns may not be directly interpreted as indications of transmissions. Our 56 sequences are matched by only 31 contemporaneous global sequences. Thus, even without any epidemiological link, Swiss sequences will cluster as they form the majority of the data. On the other hand, the high genetic similarity as well as the close sampling times and geographic origins may suggest some localized transmission. Further global sequencing efforts will be required to properly characterize local spread.

The hypervariable region between the M and F genes was unevenly recovered by our method. In some samples, the region of poor coverage extended into the beginning of the F gene, leaving the first 20 basepairs of gene F with insufficient coverage for reliable base-calling. This resulted in unknown amino acids at positions 8 and 14 of the F protein, potentially affecting the precision of their exact placement in the phylogeny. This is particularly relevant because, while previous HPIV-3 genomic studies have focused on phylogenies based on single genes like F or HN [2,51,62,63], full-length genome phylogenies are considerably more informative and can differ from these single-gene trees [64]. Interestingly, a phylogeny based on the hypervariable region itself was found to most closely resemble the full-length genome tree [65], highlighting the potential importance of this region for accurate phylogenetic inference. Given the limited knowledge about HPIV-3 infections, including the full spectrum of seasonal circulation patterns beyond the reported late Spring circulation [34,66], and ongoing disagreements about its recombination potential [67,68], our decision to utilize full-length genomes aimed to capture as much genomic data as possible.

Considering the limitations in available data and potential sampling biases, a more detailed investigation into the circulation of HPIV-3 is warranted, especially given the significant vulnerability of children to this virus and its estimated 6% global contribution to pediatric acute lower respiratory infections (ALRI) [33,34].

### The 2023/24 Swiss Influenza A/H1N1 wave seemed to be driven by multiple introductions and a single introduction expansion

Employing only open and publicly available Influenza A/H1N1 HA gene data, we reconstructed the HA phylogeny and found that the 110 Swiss samples clustered in two clades: 5a.2a and 5a.2a.1. Interestingly, based on global subsampling, irrespective of clade assignment, 71.4% (875/1225) of recent samples were from clade 5a.2a.1, while we found that 66% (101/153) of Swiss samples belonged to clade 5a.2a. In this high-density of samples within clade 5a.2a, we observed a single-variant expansion forming a Swiss monophyletic subclade (highlighted in Fig S5) as well as samples of Swiss and international origin (such as UK, US, Germany etc. found on proximal branches. The pattern is similar to a previously-described multi-introduction A/H3N2 Flu season expansion in Basel over the 2016/2017 Winter season [2] and suggests both a local expansion as well as cross-border transmissions. We find evidence for this given the proximal location of German, English and North American tips to the Swiss samples, and the progressive transmission of a single variant throughout the wave from January 19, 2024 to March 13, 2024, observed in the dense ladder-like structure of the Swiss samples, indicative of a lineage spreading over time.

### Limitations and future directions

The respiratory virus genomic surveillance framework presented in this study has several limitations. First, while hybrid capture protocols are useful to characterize viruses in parallel, including those with unknown or divergent sequences, virus identification in samples with low concentrations of viral genomic material is limited by probe design and the amount of non-viral DNA and RNA plaguing clinical samples [69,70]. Although studies have shown that full-length viral genomes can be recovered using hybrid capture protocols in samples with a CT < 30 and viruses undetected by PCR can be found [69,71], there is still uneven coverage throughout the genomes and variable coverage between different taxa [23,72,73]. Furthermore, other studies employing the same Illumina hybrid capture method found preferential coverage of certain viruses (SARS-CoV-2, RSV A/B, and Influenza A/B) at high to mid CT values (< 25), but reported dropout in other taxa, especially in samples with CT values > 30 [23,73,74]. In particular, less sensitivity to rhino/enterovirus genomes was reported, consistent with the competing TWIST panel [75]. We hypothesize that this is due to the high genetic diversity of the rhino/enterovirus families exceeding the capture efficiency of the probes employed, with up to 156 different genotypes estimated to co-circulate within a single city [76]. This uneven coverage and variability across taxa recovery biases downstream analysis, including phylogeny reconstruction, as observed with the low detection of rhino/enteroviruses from PCR-positive samples in this study (see Table S2). The protocol could be enhanced by enriching, modifying, or updating probes included in the panel to more closely capture the updated circulating viruses.

We observed a large discrepancy in assigned co-infection by sequencing in comparison to the results of PCR-testing. As some samples were hampered by low amounts of genetic material and a large by-catch of non-viral reads following targeted enrichment, it is possible that a deeper sequencing depth could have led to positive detection by the bioinformatics pipeline. Moreover, in the bioinformatics pipeline, we were conservative in (1) assigning reads to specific viruses and (2) determining the threshold indicating the presence or absence of a virus. The observed discrepancy, we hypothesize, could be attributed to: (1) the limited references present in our database in proportion to the high diversity of certain taxa and (2) discarding multi-mapper reads mapping to more than one virus in the bioinformatics pipeline. Namely, we found that HMPV genomes from the tail ends of the collection period (before 2024-02-29 and after 2024-05-21) had a dropout coverage in isolated parts of the genome compared to the other collected HMPV samples. Although these samples clustered to the B clade in the HMPV phylogeny, our bioinformatics pipeline could not distinguish between these two subtypes. The high similarity (84.38% identical) between the two genomes did not allow us to separate reads mapping to HMPV-A (accession NC_039199.1) from those mapping to HMPV-B (accession AY525843.1), even after targeting the region specific to the HMPV-A genome (5879-7152 bp). In sum, further optimizations should be carried out, particularly for taxa containing a large diversity of genomes, such as rhino/enteroviruses, to improve full-length genome recovery.

Finally, the sampling structure of our study did not allow us to discern between inpatient and outpatient samples. As such, we could not eliminate the possibility of hospital transmission influencing the observed clustering patterns, particularly in the regrouped cases observed within a single canton. Further, we lacked sufficient amounts of retrospective Swiss sequences for a formal temporal evolutionary analysis.

In this study, we constructed phylogenies using an ML frequentist approach, allowing us to rapidly assess simultaneously-circulating viruses within a single season. In the context of public health surveillance, our frequentist approach is scalable to inform at a weekly or shorter resolution, depending on the sequencing frequency, with a rapid turnaround from sampling to analysis on *multiple* co-circulating viruses. The interactive visualizations developed through Nextstrain [24] further facilitate the sharing of results. However, the framework employed does not quantify model uncertainties, as these are computationally more expensive to estimate on larger datasets, including those from this study. A promising new direction for rapid phylogenetic reconstruction incorporating Bayesian inference and uncertainty calculations is Delphy, which reduces Bayesian calculations to within a day for large datasets [77]. A combination of these tools might prove powerful for rapid viral surveillance.

In subsequent studies, once more representative global genetic diversity can be obtained, it will be possible to run more complex models analysing phylogeographical spread, evolution over multiple seasons, and epidemiological parameters inference through time. Moreover, integrating other means of surveillance in the model, such as wastewater surveillance [78], will reduce sampling bias, allowing for a complementary representative assessment of the circulating viruses. Future directions include estimating epidemiological parameters of co-circulating respiratory viruses and contextualizing this in a global context.

## Conclusion

In this proof-of-concept study, we demonstrate that time-resolved ML phylogenetic trees can be obtained from full-length genomes assembled from routine clinical samples using a hybrid capture approach, which provides additional insights on co-circulating viruses in a single country. Employing a hybrid capture approach allows us to assess a greater respiratory virus diversity than that obtained in routine multiplex PCR-testing, albeit with lower sensitivity for certain virus taxa. The workflow assembled in this study is easily adaptable to the surveillance programs of other countries, as the kit employed is off-the-shelf and the analysis pipeline is publicly available. Additional insights, particularly those regarding evolutionary relationships and mutations of interest, could not be deduced from a PCR-only strategy, highlighting the need to extend existing genomic surveillance to other viral pathogens of interest in multi-seasonal studies, which hybrid-capture panels are well-suited to capture.

## Acknowledgments

We thank Anya Parker and Chaoran Chen for setting up the Loculus instance for Influenza segments used in this study and providing a streamlined process to retrieve publicly-available SARS-CoV-2 aligned sequences. We further thank Richard Neher and Cornelius Roemer for the valuable insights on influenza clades and subclades evolutions and feedback on Nextstrain trees.

## Declaration of generative AI and AI-assisted technologies in the writing process

During the preparation of this work the author(s) used Writefull (Overleaf) and Gemini in order to proofread the manuscript. After using these tools, the author(s) reviewed and edited the content as needed and take(s) full responsibility for the content of the publication.

## Ethics approval

Sequencing of diagnostic clinical samples was approved by the Ethikkomission Nordwest- und ZentralSchweiz (EKNZ), which waived ethical approval as only viral material is processed.

## Data availability statement

The consensus sequences and raw reads from this study were deposited on ENA under Bioproject PRJNA73055. The code developed for this study can be found at https://github.com/ETH-NEXUS/2023_cb_stadler_ReVSeq and https://github.com/charlynebuerki/swiss_co-circulating_viruses. Dehumanized raw reads of samples for which we could not generate a high-quality full genome are available upon request.

