## Supplementary material for "Characterizing Co-Circulating Respiratory Virus Genomic Diversity in Switzerland with Hybrid-Capture Sequencing and Phylogenetic Reconstructions: Insights into the 2023/24 Season"

### Project Database and UI

The storage and querying of results generated by the bioinformatics analysis pipeline are facilitated through a custom web application specifically developed for this project: https://revseq.nexus.ethz.ch/ . We implemented a modular microservice architecture, where each service operates independently and agnostically, orchestrated using Docker Compose. The API backend is powered by the Django framework ( https://www.djangoproject.com/ ), while the user interface leverages Vue.js 3 (https://vuejs.org/) integrated with the Quasar framework ( https://quasar.dev/ ). Data persistence is managed through a PostgreSQL database.

Additionally, we developed a dedicated open source microservice responsible for uploading analysis results directly to the ENA Portal available at https://github.com/ETH-NEXUS/ena_upload_ms. and on DockerHub at https://hub.docker.com/r/ethnexus/ena-upload-ms.

The application offers multiple modes of interacting with the database, including searching, filtering, uploading and downloading results. Users can access data via three distinct interfaces:

1. A Swagger-based API interface, conveniently accessible from within the UI web page.
2. A command-line interface (CLI), allowing retrieval or upload of data through a set of Django management commands executed within the API container.
3. A user-friendly web application interface, enabling users to explore and filter results based on parameters such as plate or substrain.

This multi-interface design ensures flexibility, catering to various user preferences and enhancing the overall accessibility and usability of the system.

### Other virus contextual dataset assembly

For all other viruses (4 seasonal coronaviruses, HPIV-1,2, Adenovirus B1 and C2), contextual background sequences were gathered by querying publicly available consensus sequences from NCBI GenBank. These were queried using the relevant associated taxon id for each virus (HPIV-1: 12730, HPIV2: 2560525, cov-229E: 11137, cov-HKU1: 290028, cov-NL63: 277944, cov-OC43: 31631, adv-B1: 108098, adv-C2: 129951). The queried sequences were aligned and processed with Nextclade [25], where a coverage score was calculated for each consensus genome based on the same reference genome used to assemble the study's core samples (HPIV-1: NC_003461, HPIV-2: NC_003443, cov229E: NC_002645, cov-HKU1: NC_006577, cov-NL63:

NC_005831, cov-OC43: AY391777, adv-B1: NC_011203,adv-C2: NC_001405). We filtered out background sequences that had 40% or more of unknown nucleotides from the whole genome or if the Nextstrain quality-control score was flagged as "bad". For viruses with large collections of sequences, We further subsampled sequences to 100 sequences per country per year and removed any sequences sampled before 1925.

For HMPV, we modified the existing Nextstrain build (https://github.com/nextstrain/hmpv) to incorporate this study’s core samples and required at least 80% full-genome coverage.

For SARS-CoV-2 contextual background sequences, we queried the LAPIS API [26] to obtain all publicly available SARS-CoV-2 aligned consensus sequences and filtered sequences to include European sequences and a subset of other representative sequences from around the world, stemming from the Omicron variant of concern (all sequences used are available here: https://nextstrain.org/ncov/open/europe/all-time?label=clade:21M%20%28Omicron%29). We used the strain "Wuhan/IPBCAMS-WH-02/2019" (accession: MT019530.1) to root the phylogeny. We employed the Nextstrain classic clade naming scheme.

For Influenza A/H3N2 and Influenza B/Victoria we proceeded similarly to A/H1N1, where we queried GenSpectrum ( https://loculus.genspectrum.org/) to obtain all publicly available consensus sequences subtyped as H3 and N2 and Victoria, subsequently downsampling sequences to 10 sequences per year from when the virus emerged (2007-01-01 for A/H3N2 and 2008-01-01 for B/Victoria) until 2022, and 300 sequences per country per year for the last 2 years. NA and HA segments were aligned and sequences with segments containing more than 5% of unknown nucleotides were dropped. The phylogenies were rooted to their respective pandemic strains (H3N2 HA segment: CY163680.1, B/Victoria HA segment: CY115151.1). We constructed trees for segment 4 (HA) and segment 6 (NA) and employed classic clade naming schemes available on Nextstrain to label clades (https://github.com/ influenza-clade-nomenclature).

All builds are available on the project github and can be visualized here: https://nextstrain.org/community/cevo-public/ReVSeq-project

### Supplementary Tables

Table S1: Counts of all genomes included in the phylogenetic trees by strain.

| Strain | Total no. genomes | No. context genomes | No. Study |
|---|---|---|---|
| **RSV-A** | 3074 | 3018 | 56 |
| **RSV-B** | 2643 | 2632 | 11 |
| **HPIV-3** | 823 | 767 | 56 |
| **Influenza A/H1N1 HA** | 1607 | 1497 | 110 |
| **Influenza A/H1N1 NA** | 1727 | 1613 | 114 |
| **HMPV** | 1015 | 986 | 29 |
| **HPIV-1** | 234 | 231 | 3 |
| **HPIV-2** | 102 | 100 | 2 |
| **Influenza A/H3N2 HA** | 2860 | 2848 | 12 |
| **Influenza A/H3N2 NA** | 2848 | 2838 | 10 |
| **Influenza B/Victoria HA** | 1137 | 1110 | 27 |
| **Influenza B/Victoria NA** | 1147 | 1120 | 27 |
| **SARS-CoV-2** | 1048 | 987 | 61 |
| **HKU1** | 271 | 264 | 7 |
| **OC43** | 526 | 514 | 12 |
| **NL63** | 282 | 278 | 4 |
| **229E** | 281 | 276 | 5 |
| **Adenovirus B1** | 752 | 735 | 17 |
| **Adenovirus C2** | 437 | 428 | 9 |

Table S2: Counts of virus strains identified in each type of testing employed (PCR positivity, detection by sequencing and high-quality genomes extracted. Grey squares indicate that the testing employed cannot detect these strains.

| | | PCR panel | | | Sequencing | |
|---|---|---|---|---|---|---|
| **family** | **Strain** | Resp | RSV/ Inf/SARS | Inf/RSV | Detection | HQ |
| paramyxoviridae | RSV-A/B | 61 | 17 | 35 | 144 | 67 |
| | HMPV | 82 | | | 62 | 29 |
| | HPIV-1 | 5 | | | 8 | 3 |
| | HPIV-2 | 4 | | | 4 | 2 |
| | HPIV-3 | 88 | | | 72 | 56 |
| | HPIV-4 | 3 | | | 2 | 2 |
| Orthomyx oviridae | Influenza A | 89 | 74 | 42 | 177 | 131 |
| | Influenza B | 15 | 12 | 15 | 65 | 28 |
| coronaviridae | SARS-CoV-2 | 68 | 52 | | 125 | 61 |
| | HKU1 | 23 | | | 14 | 7 |
| | OC43 | 31 | | | 28 | 12 |
| | NL63 | 11 | | | 9 | 4 |
| | 229E | 14 | | | 18 | 5 |
| adenovi ridae | Adenovirus | 84 | | | 148 | 28 |
| others | Rhino/enterovirus | 322 | | | 99 | 12 |
| | Parechiovirus | | | | 4 | 1 |
| | bocavirus | | | | 6 | 5 |
| | Polyomavirus | | | | 15 | 8 |

Table S3: $R^2$ correlations between the PCR-positive strains and detected strains by sequencing from this study with the 2 national surveillance systems: Sentinella and wastewater.

| | **Study samples** | | **Sentinella [%positive PCR]** | | **Wastewater viral load [gc/person/day]** | |
|---|---|---|---|---|---|---|
| **Strain** | PCR / seq. detected | Seq. detected / seq. hq | PCR / %pos. | Seq. detected / %pos | PCR / viral load | Seq detected / viral load |
| Influenza A | 0.94 | 0.93 | 0.65 | 0.60 | 0.75 | 0.71 |
| Influenza B | 0.72 | 0.64 | 0.20 | 0.21 | 0.27 | 0.22 |
| RSV-A/B | 0.53 | 0.92 | 0.11 | 0.09 | 0.52 | 0.32 |
| SARS-CoV-2 | 0.68 | 0.62 | 0.18 | 0.15 | 0.38 | 0.32 |
| Adenovirus | 0.58 | 0.72 | 0.01 | 0.02 | | |
| Rhino/enterovirus | 0.22 | 0.03 | 0.03 | 0.0 | | |
| others | 0.77 | 0.72 | 0.08 | 0.02 | | |

Table S4: Number of full genome sequences publicly available on NCBI and from Switzerland for a selection of viruses covered in this study, as of March 31, 2025.

| **Strain** | **Taxon ID** | **No. NCBI** | **No. Switzerland** | **No. study** |
|---|---|---|---|---|
| **RSV-A** | 208893 | 5055 | 0 | 56 |
| **RSV-B** | 208895 | 4293 | 1 | 11 |
| **HPIV-3** | 11216 | 804 | 1 | 56 |
| **HMPV** | 162145 | 1078 | 0 | 29 |
| **Influenza H1N1(HA)** | 114727 | 22277 | 54 | 118 |
| **HPIV-1** | 12730 | 247 | 1 | 3 |
| **HPIV-2** | 2560525 | 144 | 0 | 2 |
| **OC43** | 31631 | 568 | 0 | 12 |
| **229E** | 11137 | 314 | 0 | 5 |
| **HKU1** | 290028 | 327 | 0 | 7 |
| **NL63** | 277944 | 310 | 0 | 4 |

### Supplementary Figures

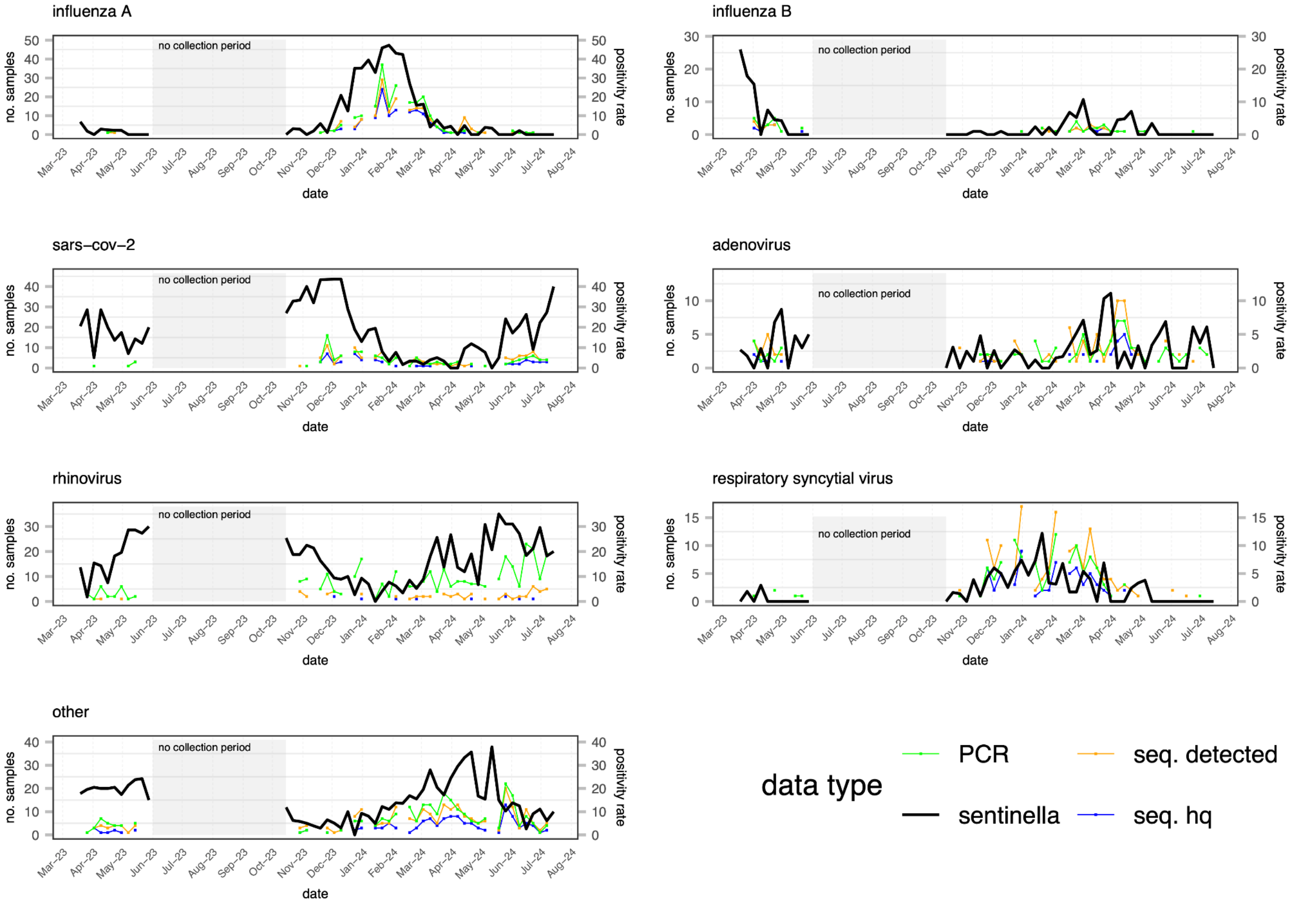


Figure S1: **Time series of the counts of this study's samples and the national Sentinella surveillance by pathogen at a weekly resolution**: each panel plots the weekly positivity rate from the Sentinella data (black), the number of PCR-positive samples collected (green) the number of detected strains (orange), and the number of high-quality sequences generated (blue).

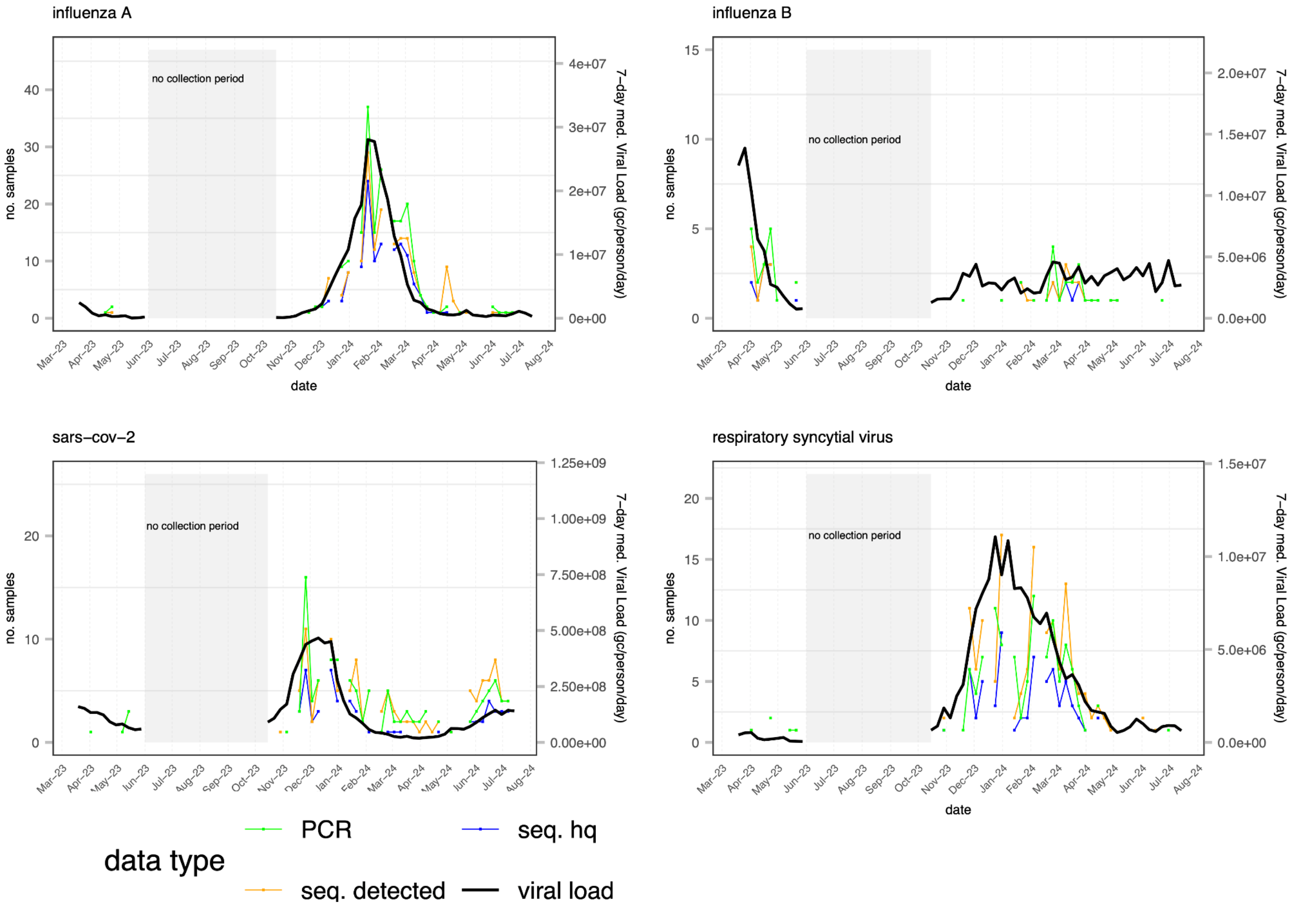


Figure S2: **Time series of the counts of this study's samples and the national wastewater surveillance by pathogen at a weekly resolution**: each panel plots the 7-day median viral load [gc/person/day] reported from the national wastewater surveillance (black), the number of PCR-positive samples collected (green) the number of detected strains (orange), and the number of high-quality sequences generated (blue).

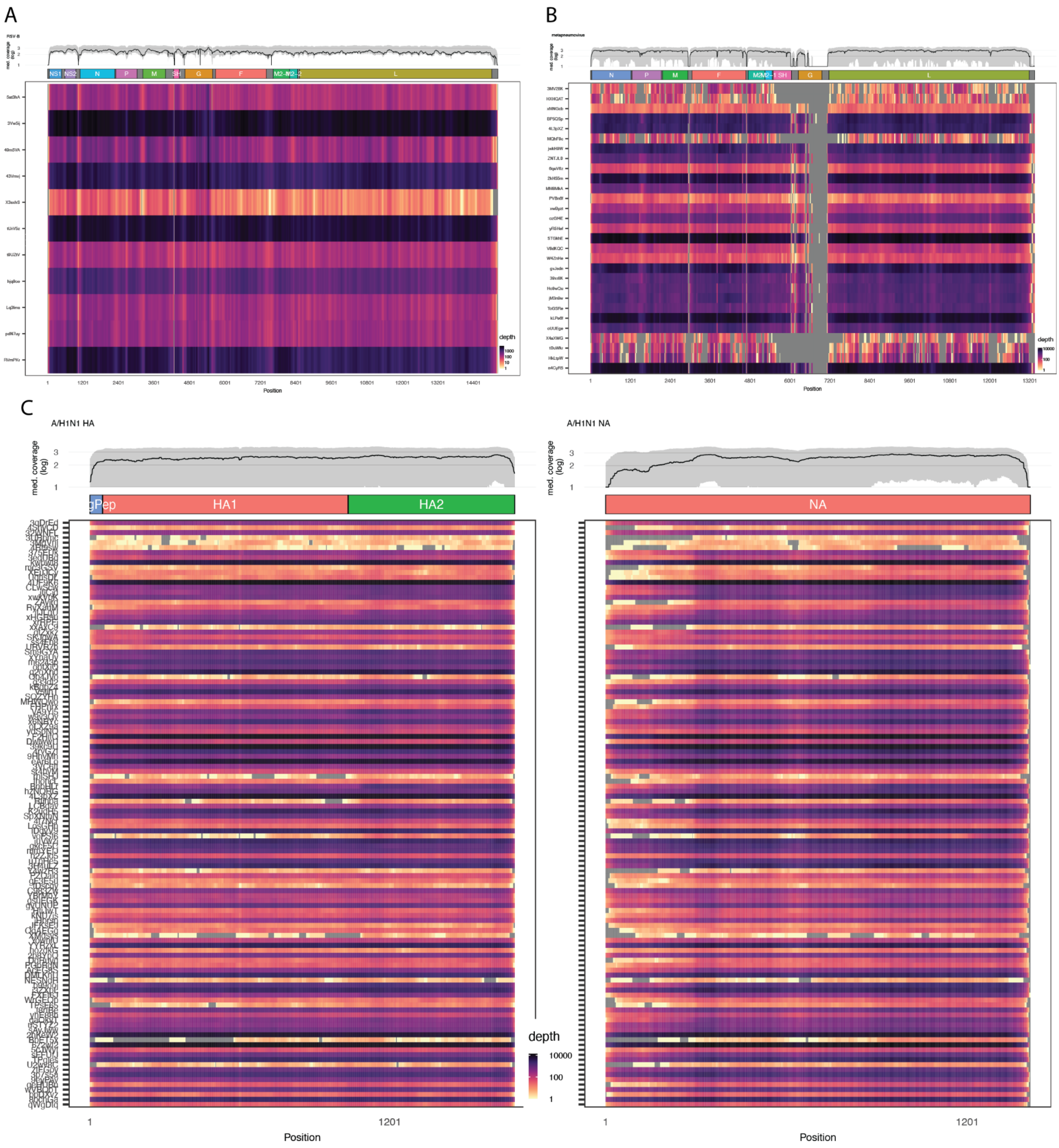


Figure S3: **Whole-genome coverage of RSV-B, HMPV and H1N1 HA and NA segments**: The 11 RSV-B samples in **A.**, 29 HMPV samples in **B.**, and 118 influenza A/H1N1 samples in **C.** annotated, whole-genome coverage of high-quality genomes obtained in the study. The median coverage in black and the $10^{th}$ to $90^{th}$ quartile in shaded grey calculated at each basepair is plotted in the top panel, and the individual genomes's depths are plotted as a heatmap below, where a darker color indicates greater depth. Grey indicates a depth coverage of 0. Samples are ordered in increasing collection date order.

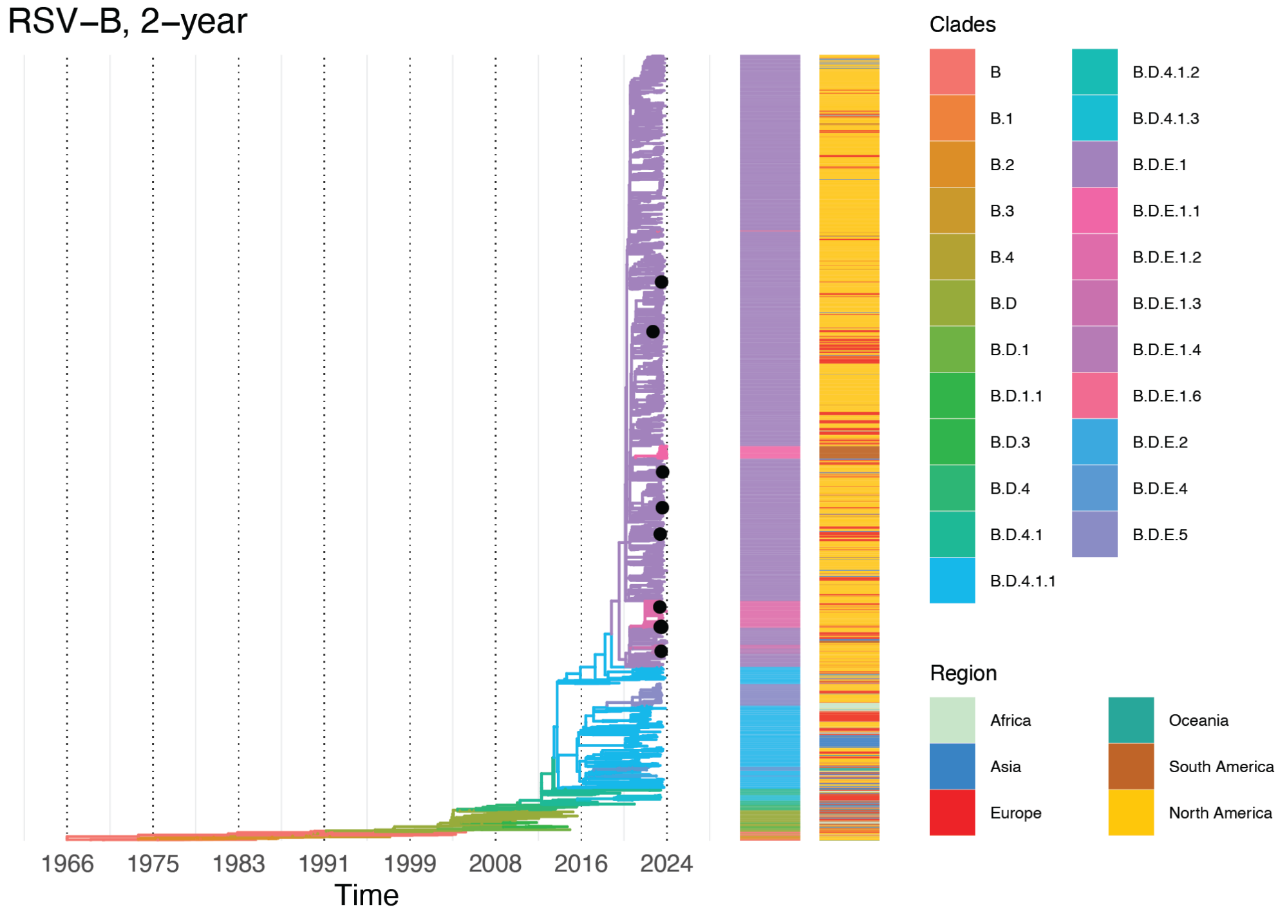


Figure S4: **whole-genome phylogeny of RSV-B** The maximum likelihood phylogenetic time tree generated with the Nextstrain build color-coded by clade and labelled with black dots for this study's samples (N=12). Additional geographical information is annotated to the right of the tree. The clade labeling is from Nextstrain.

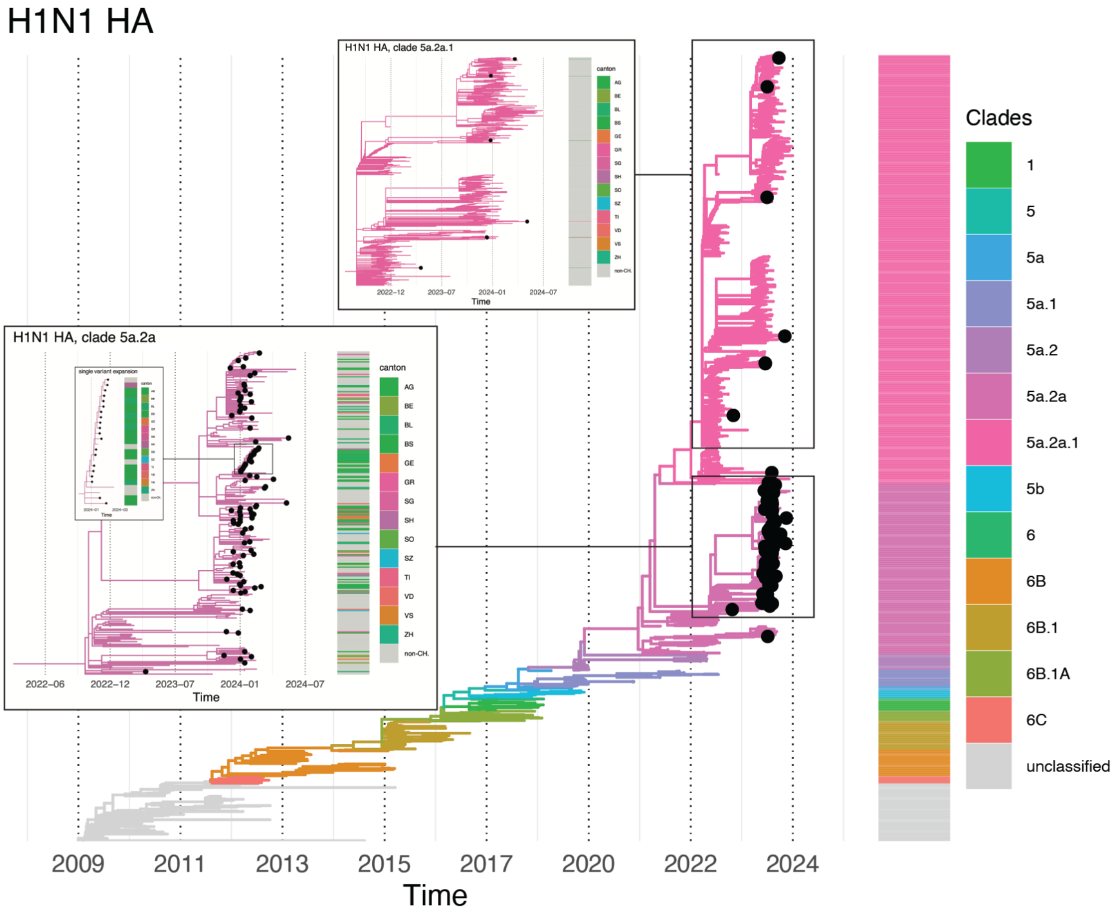


Figure S5: **HA phylogeny of Influenza A/H1N1** The maximum likelihood phylogenetic time tree generated with the Nextstrain build color-coded by clade and labelled with black dots for this study's samples (N=110) with 2 insets into clade 5a.2a.1 (top; pink) and clade 5a.2 (bottom; purple), with the cantonal location of this study's samples to the right of the insets. The clade labeling is from Nextstrain. In inset 5a.2a, we include a zoom into the single variant expansion found in this study's samples.
